# The ApoC2 mimetic peptide D6PV enhances remyelination by stimulating oxidative phosphorylation in oligodendrocytes

**DOI:** 10.64898/2026.01.22.701034

**Authors:** Brecht Moonen, Laura Bolkaerts, Kirsten Poelmans, Flore Wouters, Koen Kuipers, Jeroen Guns, Sanne G.S. Verberk, Esther Wolfs, Yvonne Jansen, Yvonne Döring, Tim Vanmierlo, Tess Dierckx, Erin Gibson, Anna Wolska, Mart Reimund, Alan T. Remaley, Melanie Loix, Jerome J.A. Hendriks, Jeroen F.J. Bogie, Sam Vanherle

## Abstract

Failure of remyelination drives neurodegeneration in demyelinating disorders such as multiple sclerosis (MS), with disrupted lipid handling and metabolic stress in oligodendrocyte precursor cells (OPCs) posing major barriers to repair. Here, we identify the dual ApoC-II mimetic–ApoC-III antagonist peptide D6PV as a metabolic modulator that directly enhances OPC differentiation and myelin repair. Across *ex vivo* and *in vivo* models of chemically induced demyelination, D6PV promotes oligodendrocyte maturation and restores myelin integrity independently of lipoprotein hydrolysis or modulation of lipid droplet-containing phagocytes. Guided by transcriptomics analyses, we find that D6PV stimulates mitochondrial oxidative phosphorylation and fatty acid β-oxidation, while suppressing inflammatory transcriptional programs, thereby driving OPCs toward a myelinating phenotype. Notably, D6PV does not alter peripheral immune composition or autoimmune-driven pathology in the experimental autoimmune encephalomyelitis model, indicating a central nervous system (CNS) cell-autonomous effect. These findings reveal a metabolism-linked pathway for remyelination and position D6PV as a promising therapeutic strategy to enhance CNS repair in demyelinating diseases.

## Introduction

Central nervous system (CNS) demyelination, arising from the loss or damage of myelin sheaths, is a defining feature of numerous neurological disorders and a major driver of axonal degeneration and functional decline ^1–4^. Efficient remyelination is essential for restoring neural function; however, in chronic disease stages in CNS disorders, this regenerative process often falters ^1,2,5,6^. Multiple sclerosis (MS), the most prevalent demyelinating disorder, exemplifies this challenge. Despite oligodendrocyte precursor cells (OPCs) being recruited to demyelinated lesions, their differentiation and myelin-forming capacity are frequently impaired, leading to the formation of chronically demyelinated lesions ^5–7^. The mechanisms underlying this failure remain incompletely understood, yet accumulating evidence implicates both intrinsic deficits in OPCs and extrinsic constraints imposed by the lesion microenvironment ^8–16^. Notably, disruptions in intercellular lipid handling and lipoprotein-mediated lipid transfer have emerged as key determinants of this impaired regenerative capacity ^14^.

Increasing evidence indicates that key regulators of triglyceride metabolism, including apolipoproteins C-II (ApoC-II) and C-III (ApoC-III), as well as their shared enzymatic target lipoprotein lipase (LPL), are dysregulated in MS ^17–19^. ApoC-II serves as an essential activator of LPL, promoting the hydrolysis and clearance of triglyceride-rich lipoproteins ^20^, whereas ApoC-III antagonizes LPL activity and delays lipoprotein catabolism, thereby favoring triglyceride accumulation in lipoproteins ^21^. Transcriptomic and proteomic analyses have reported altered expression of LPL and its regulatory apolipoproteins in MS lesions and circulating immune cells, consistent with impaired triglyceride processing and abnormal lipid handling ^17,22^. Functionally, lipid profiles characteristic of ApoC-II/ApoC-III imbalance - including elevated circulating triglycerides - correlate with disease activity and disability in MS, supporting a link between disrupted triglyceride metabolism, neuroinflammation, and disease progression ^23–26^. Despite this, whether targeting ApoC-II/ApoC-III–LPL signaling can modulate neuroinflammation or enhance regenerative responses in the CNS remains unknown.

Here, we examined the effects of the dual ApoC-II mimetic/ApoC-III antagonist peptide D6PV across complementary models of autoimmune- and chemically-induced demyelination. We find that D6PV does not alter autoimmune-driven pathology in the experimental autoimmune encephalomyelitis (EAE) model, and immune phenotyping confirms no detectable changes in the abundance or composition of major immune cell subsets within peripheral lymphoid organs. In contrast, D6PV markedly enhances remyelination in cerebellar brain slice cultures and the cuprizone model by promoting OPC differentiation, an effect that appears largely independent of its modulation of lipoproteins. Mechanistically, we show that D6PV enhances mitochondrial oxidative phosphorylation in OPCs, thereby facilitating their maturation. Together, these findings identify a previously unrecognized, metabolism-linked mechanism to promote CNS regeneration, and position D6PV as a promising therapeutic strategy for enhancing remyelination.

## Methodology

### Antibodies and chemical reagents

The following antibodies were used for immunofluorescent/immunohistochemical stains: rabbit anti-NF (1:1000, Ab8135, Abcam), rat anti-MBP (1:250, MCA409S, Millipore, *ex vivo* cerebellar brain slices), rat anti-MBP (1:500, MAB386, Millipore, brain cryosections and *in vitro* OPC cultures), mouse anti-CC1 (1:50, ab16794, Abcam), goat anti-olig2 (1:50, AF2418, R&D systems), mouse anti-O4 (1:1000, MAB1326, R&D systems). Appropriate secondary antibodies were purchased from Invitrogen. ApoA-I (50 µg/mL, kindly provided by Alan Remaley, National Health Institute, Bethesda, USA) was used as a cholesterol acceptor in the cholesterol efflux assay.

### Peptide D6PV

Peptide D6PV was kindly provided by Alan Remaley (National institute of Health). Briefly, peptide D6PV was synthesized using standard Fmoc chemistry by Wuxi AppTech (China) and purified to greater than 95% purity by reverse-phase high-performance liquid chromatography on an Aquapore RP-300 column.

### Animals

Wild-type C57BL/6J mice were purchased from Envigo and were housed in a 12h light/dark cycle with ad libitum access to water and a standard chow diet or specific formulations as indicated. All animal procedures were conducted in accordance with the institutional guidelines and approved by the Ethical Committee for Animal Experiments of Hasselt University (protocol numbers 202403B, 202485, 202486).

### Cerebellar brain slice cultures

Cerebellar brain slices were obtained from wild-type C57BL/6J mouse pups at postnatal day 10 (P10), as described previously ^27,28^. Brain slices were cultured in MEM medium (Thermo Fisher Scientific) supplemented with 25% horse serum (Thermo Fisher Scientific), 25% Hank’s balanced salt solution (Sigma-Aldrich), 50 U/mL penicillin (Invitrogen), 50 U/mL streptomycin (Invitrogen), 1% Glutamax (Thermo Fisher Scientific), 12.5 mM HEPES (Thermo Fisher Scientific), and 1.45 g/L glucose (Sigma-Aldrich) at 37°C and 5% CO_2_. For phagocyte depletion, slices were treated with clodronate or empty liposomes (0.5 mg/mL, LIPOSOMA) immediately after isolation for 24h. To induce demyelination, brain slices were treated with 0.5 mg/mL lysolecithin (Sigma-Aldrich) at 3 days post-isolation for 16 h. Next, brain slices were allowed to recover in culture medium for 1 day and subsequently treated daily for 6 days with vehicle or D6PV (50 µM), followed by histological and biochemical analyses.

### Cuprizone-induced acute demyelination *in vivo* model

To induce acute demyelination, 9-week-old male wild-type mice were fed *ad libitum* a diet containing 0.3% (w/w) cuprizone (bis[cyclohexanone]oxaldihydrazone, Sigma-Aldrich) mixed in powdered standard chow for 5 weeks. Subsequently, cuprizone diet was withdrawn to allow spontaneous remyelination for 1 week. During this week, mice were treated daily intraperitoneally with D6PV (23.4 mg/kg body weight ^29^). After 5 weeks or 6 weeks from the start of cuprizone diet, mice were sacrificed, and tissue was collected for histological and biochemical analyses.

### Experimental autoimmune encephalomyelitis

11-week-old female mice were immunized subcutaneously with myelin oligodendrocyte glycoprotein peptide 35–55 (MOG_35–55_) emulsified in complete Freund’s adjuvant with Mycobacterium tuberculosis according to the manufacturer’s guidelines (EK-2110 kit; Hooke Laboratories, United States, Lawrence). Directly after immunization and after 24 h, the mice were intraperitoneally (IP) injected with 110 ng pertussis toxin in PBS. Starting 5 days post-immunization, the mice were treated daily IP with D6PV (23.4 mg/kg) or vehicle (20 mM Tris). Mice were weighed daily and clinically evaluated for neurological signs of the disease in a blinded fashion following a five-point standardized rating of clinical symptoms (0: no clinical symptoms; 1: tail paralysis; 2: tail paralysis and partial hind limb paralysis; 3: complete hind limb paralysis; 4: paralysis to the diaphragm; 5: death by EAE). Spleen and lymph nodes were isolated at 15 days post-immunization (dpi, peak) and after 27 dpi (chronic).

### Plasma lipid and lipoprotein analysis

Cholesterol and triglyceride levels were analyzed using mouse EDTA-buffered plasma and quantified using enzymatic assays (c.f.a.s. cobas, Roche Diagnostics) according to the manufacturer’s instructions. Plasma very low-density lipoproteins (VLDL), low-density lipoproteins (LDL), and high-density lipoproteins (HDL) were separated by fast performance liquid chromatography (FPLC)-size exclusion chromatography analysis (gel filtration on a Superose 6 column (GE Healthcare) at a flow rate of 0.5 mL/min). The different lipoprotein fractions were collected according to their retention times as follows: VLDL between 40 and 50 min, LDL between 50 and 70 min, and HDL between 70 and 90 min. Results are presented as optical density (OD) profiles at 492 nm of the isolated lipoproteins.

### Myelin isolation

Myelin was purified from post-mortem mouse brain tissue of male and female C57BL6/J mice by means of density gradient centrifugation, as described previously ^30^. Briefly, brain tissue was homogenized in 0.32M sucrose and subsequently layered on top of 0.85M sucrose. After centrifugation at 75,000g, myelin was collected from the interface and washed and purified in water. Myelin protein concentration was determined using the BCA protein assay kit (Thermo Fisher Scientific), according to the manufacturer’s guidelines.

### Cell culture (BMDMs and OPCs)

Bone marrow-derived macrophages (BMDMs) were obtained as described previously ^30^. Briefly, femoral and tibial bone marrow was isolated from 11-week-old female wild-type C57BL/6J mice and cultured in 14.5 cm petri dishes (Greiner, ref. 639161) at a concentration of 10 × 10^6^ cells/petri dish in RPMI1640 medium (Lonza) supplemented with 10% fetal calf serum (FCS; Biowest), 50 U/mL penicillin (Invitrogen), 50 U/mL streptomycin (Invitrogen), and 15% L929-conditioned medium (LCM) for 7 days at 37°C and 5% CO_2_. Next, cells were cultured at a density of 0.5 × 10^6^ cells/mL in RPMI1640 supplemented with 10% FCS, 50 U/mL penicillin, 50 U/mL streptomycin, and 5% LCM. BMDMs were treated daily with vehicle, 10 µM D6PV and/or 100 µg/mL myelin. For phenotypic analysis, BMDMs were stimulated with LPS and IFN-γ (both 100 ng/mL) for 6h to assess gene expression.

OPCs were isolated from P0-2 wild-type C57BL/6J mice cerebral cortices, and cells were enzymatically dissociated for 20 min at 37°C with papain (3 U/mL, Sigma-Aldrich), diluted in Dulbecco’s Modified Eagle Medium (DMEM; Gibco) and supplemented with 1 mM L-cysteine (Sigma-Aldrich), and DNase I (20 µg/mL, Sigma-Aldrich) using the gentleMACS^TM^ dissociator (Miltenyi Biotec). The resulting cell suspension was passed through a 70 µm strainer to obtain a single-cell suspension. OPCs were subsequently isolated by magnetic-activated cell sorting (MACS) using the CD140a (PDGFRa) MicroBead Kit mouse (5250108305, Miltenyi Biotec) according to manufacturer’s instructions. Afterwards, OPCs were plated on poly-L-lysine (PLL)-coated wells at a density of 3 x 10^5^ cells/mL in differentiation medium [DMEM medium supplemented with 0.3 mM transferrin, 0.1 mM putrescin, 0.02 mM progesterone, 0.2 µM sodium selenite, 0.5 µM triiodothyronine, 0.5 mM L-thyroxin, 0.8 mM bovine insulin, 0.5% P/S, 2 % horse serum, 2 % B27 supplement; all from Sigma-Aldrich except for P/S (Invitrogen) and B27 (Thermofisher)] for 6 days at 37°C and 8.5% CO_2_. OPCs were treated daily with vehicle or D6PV (10 µM) for 6 days, followed by histological and biochemical analyses.

### Immunofluorescence

Murine OPCs cultured on PLL-coated glass cover slides were fixed in 4% paraformaldehyde (PFA) for 20 min at room temperature. To stain OPCs, samples were incubated with relevant primary antibodies diluted in blocking buffer (1x PBS + 1% BSA + 0.1% Tween-20). Imaging of OPCs was performed using a LeicaDM2000 LED fluorescence microscope. ImageJ was used to determine the MBP/O4-positive area per cell and to define the Sholl analysis parameters related to process complexity and branching (*i.e.*, sum intersections, average intersections/Sholl ring, end radius) in OPCs as previously described ^31^. Cerebellar brain slices were fixed in 4% PFA for 15 min at room temperature. To stain cerebellar brain slices, samples were incubated with relevant primary antibodies diluted in blocking buffer (1x PBS + 5% horse serum + 0.3% Triton X-100). Imaging was performed using an LSM880 confocal microscope (Zeiss). ImageJ was used to define the Olig2 and CC1 positive cells, which are presented as the percentage Olig2^+^ CC1^+^ cells among all Olig2^+^ cells and the myelination index (MBP^+^ NF^+^ axons/NF^+^ axons), which is presented in a relative normalized manner. Three-dimensional analysis of cerebellar brain slices was performed using the z-stack feature, and images were 3D rendered using the 3D rendering software Vaa3D ^32^.

Brain tissue of cuprizone mice was isolated, snap-frozen, and sectioned with a Leica CM1900UV cryostat (Leica Microsystems) to obtain 10 µm slices. Cryosections were fixed in ice-cold acetone for 10 min at −20°C. Immunostaining and analysis of cryosections were performed as described previously ^15,30^. Cryosections were imaged using a LeicaDM2000 LED fluorescence microscope. ImageJ was used to align the corpus callosum, followed by determination of the MBP, Olig2, and CC1 positive signal in this area.

### Transmission electron microscopy

Mouse brain samples were fixed with 2% glutaraldehyde. Next, post-fixation was done with 2% osmiumtetroxide in 0.05 M sodium cacodylate buffer for 1h at 4°C. Dehydration of the samples was performed by ascending concentrations of acetone. Next, the dehydrated samples were impregnated overnight in a 1:1 mixture of acetone and araldite epoxy resin. Next, the samples were embedded in araldite epoxy resin at 60°C and cut into slices of 70 nm, perpendicular to the corpus callosum, with a Leica EM UC6 microtome. The slices were transferred to copper grids (Veco B.V) that were coated with 0.7% formvar. Analysis was performed with a Jeol JEM-1400 Flash at 80 kV equipped with an Emsis 20NP XAROSA camera system; and around 10 images were taken of different regions in the corpus callosum. ImageJ was used to calculate the g-ratio (ratio of the inner axonal diameter to the total outer diameter). All analyses were conducted by observers blinded to the experimental arm of the study.

### Flow cytometry

For the staining of spleen- and lymph node-derived lymphocytes, the following antibodies were used (all purchased from BioLegend): CD45 Alexa Fluor 700 (clone 30-F11, 1:200), CD19 BV650 (clone 6D5, 1:200), CD11b BV785 (clone M1/70, 1:200) CD3 FITC (clone 17A2, 1:200), CD4 BV510 (clone RM4-5, 1:200), CD8 PerCP-Cy5.5 (clone 53-6.7, 1:200), IFNγ BV605 (clone XMG1.2, 1:100), IL17 PE-Dazzle 594 (clone TC1-18H10.1, 1:100), IL4 PE (clone 11B11, 1:100), and FOXP3 Alexa Fluor 647 (clone MF-14, 1:100). Dead cells were excluded by Fixable Viability Dye Zombie NIR (1:1000, BioLegend). For staining of intracellular cytokines, cells were stimulated with phorbol 12-myristate 13-acetate (20 ng/ml; Sigma‒Aldrich), calcium ionomycin (1 µg/ml; Sigma‒Aldrich), and Golgiplug (2 µg/ml; BD Biosciences) for 4 h, and the FOXP3/transcription factor staining buffer set (Invitrogen) was used according to the manufacturer’s instructions. Cells were acquired on an Aurora spectral flowcytometer (Cytek Biosciences).

### Phagocytosis assay

To evaluate the ability and extent of myelin phagocytosis, myelin was fluorescently labelled with pHrodo succinimidyl ester (cat. No. P36600, Invitrogen). Cells were exposed to 100 µg/ml pHrodo-labeled myelin for 1.5h at 37°C and analyzed for fluorescence intensity using the CLARIOstar PLUS microplate reader.

### Bodipy and Oil red O stains

BMDMs were stained for intracellular lipid load by 15 min incubation with BODIPY (493/503) at 37°C. The FACS Fortessa (BD Biosciences) was used to quantify cellular fluorescence. For Oil red O (ORO) stains, cells were fixed with 4% PFA for 15 min. Next, cells were stained with 0.3% ORO for 10 min and counterstained with nuclei stain haematoxylin for 1 min. Analysis was carried out using a Leica DM 2000 LED microscope and ImageJ software.

### Cholesterol measurements

Cholesterol levels of BMDMs were defined by using the Amplex Red Cholesterol Assay kit (Thermo Fisher) according to the manufacturer’s instructions. To determine the cholesterol efflux of OPCs, OPC cultures were exposed to peptide 5A (50 µg/mL) in phenol- and serum-free medium for 4h before measuring the intra- and extracellular total cholesterol levels. Fluorescence was measured using the CLARIOstar PLUS microplate reader (BMG Labtech), and cholesterol efflux was determined by dividing fluorescence in the supernatants by the total fluorescence in supernatants and cells.

### Quantitative PCR (qPCR)

Cells were lysed using QIAzol lysis reagent (Qiagen). Next, RNA was extracted using the RNeasy mini kit (Qiagen) according to the manufacturer’s instructions. cDNA was synthesized using the qScript cDNA synthesis kit (Quanta Biosciences) according to the manufacturer’s instructions. Quantitative PCR was performed on a StepOnePlus detection system (Applied Biosystems). Data were analyzed using the comparative Ct method and normalized to the most stable reference genes as determined by GeNorm, being *Cyca* and *Rpl13a* for RNA isolated out of cerebellar brain slices and corpus callosum tissue of cuprizone animals, and *Cyca* and *Tbp* for RNA derived from BMDMs. Primer sequences are available upon request.

### Measurements of reactive oxygen species

After treatment, BMDMs were stimulated with phorbol 12-myristate 13-acetate (PMA, 100 ng/ml, Sigma-Aldrich) for 15 min. Next, reactive oxygen species (ROS) production was measured using the fluorescent probe 2’,7’-dichlorodihydrofluorescein diacetate at 10 µM in PBS for 30 min. Fluorescence was measured using the CLARIOstar PLUS microplate reader (BMG Labtech).

### RNA sequencing

Total RNA from OPCs was isolated using QIAzol and the RNeasy mini kit, according to the manufacturer’s guidelines. Preparation of RNA libraries and sequencing was outsourced to Novogene, specifying Eukaryotic RNA-Seq as the library type, followed by sequencing on a Novaseq X Plus (Illumina) with a PE 150 strategy. Following quality control, reads were aligned to the mouse reference genome (Mus musculus, GRCm38/mm10 release M21) and transcript abundancies were quantified using Salmon (1.4.0) using the --gcBias and –seqBias flags ^33^. Gene-level count matrices were created using the tximport package (v1.18.0) and used as input for DEG analysis with DESeq2 (v1.30.1) as previously reported ^34,35^. PCA analysis was performed using the plotPCA function from the DESeq2 package after rlog transformation. Based on PCA analysis, two samples (one vehicle control, one D6PV-treated) were identified as outliers and omitted from further downstream analyses. For visualization and gene ranking, log fold change (LFC) estimates were shrunk using the apeglm (v1.12.0) method for effect size shrinkage as described by ^34,36^. Pathway analysis of differentially expressed genes was conducted using Ingenuity Pathway Analysis.

### Statistical analysis

Data were statistically analyzed using GraphPad Prism v8 and are reported as mean ± s.e.m.. The number of animals, slices, cultures, and biological replicates is indicated in the figure legends. Measurements were taken from distinct samples. Data collection was randomized for all experiments. The D’Agostino and Pearson omnibus normality test was used to test for normal distribution. When datasets were normally distributed, an ANOVA (Tukey’s post hoc analysis) or two-tailed unpaired Student’s *t*-test (with Welch’s correction if necessary) was used to determine statistical significance between groups. If the dataset did not pass normality, the Kruskal-Wallis or Mann-Whitney analysis was applied. p values <0.05 were considered to indicate a significant difference (*, p < 0.05; **, p < 0.01; ***, p < 0.001, ****, p < 0.0001).

## Results

### Peptide D6PV promotes remyelination in the cerebellar brain slice and cuprizone models

Given the central role of lipid metabolism in brain homeostasis and repair ^14^, we investigated whether the dual ApoC-II mimetic–ApoC-III antagonist peptide D6PV could impact remyelination in models of chemically induced demyelination. We first evaluated the remyelinating potential of D6PV in an *ex vivo* system using organotypic cerebellar brain slice cultures (BSCs), which preserve the native CNS architecture and allow for the study of myelin repair in a physiologically relevant setting, recapitulating key aspects of endogenous remyelination, including oligodendrocyte maturation and glial cell responses ^15,16,37^. Demyelination was induced by lysophosphatidylcholine (LPC), and slices were treated with D6PV (50 µM) during the recovery phase (experimental design depicted in **Fig. 1A**). Immunofluorescence analysis revealed a marked increase in colocalization between myelin basic protein (MBP) and neurofilament (NF) in BSCs exposed to D6PV (**Fig. 1B-C**), indicative of restored myelin coverage along axons. These effects were further visualized through three-dimensional reconstructions using Vaa3D ^32^, which showed improved myelin architecture in D6PV-treated slices. In parallel, quantification of oligodendrocyte lineage cells revealed a significant increase in the proportion of mature Olig2⁺ CC1⁺ oligodendrocytes within the total Olig2⁺ population, pointing toward D6PV promoting oligodendrocyte differentiation (**Fig. 1B,D**).

**Figure 1:**
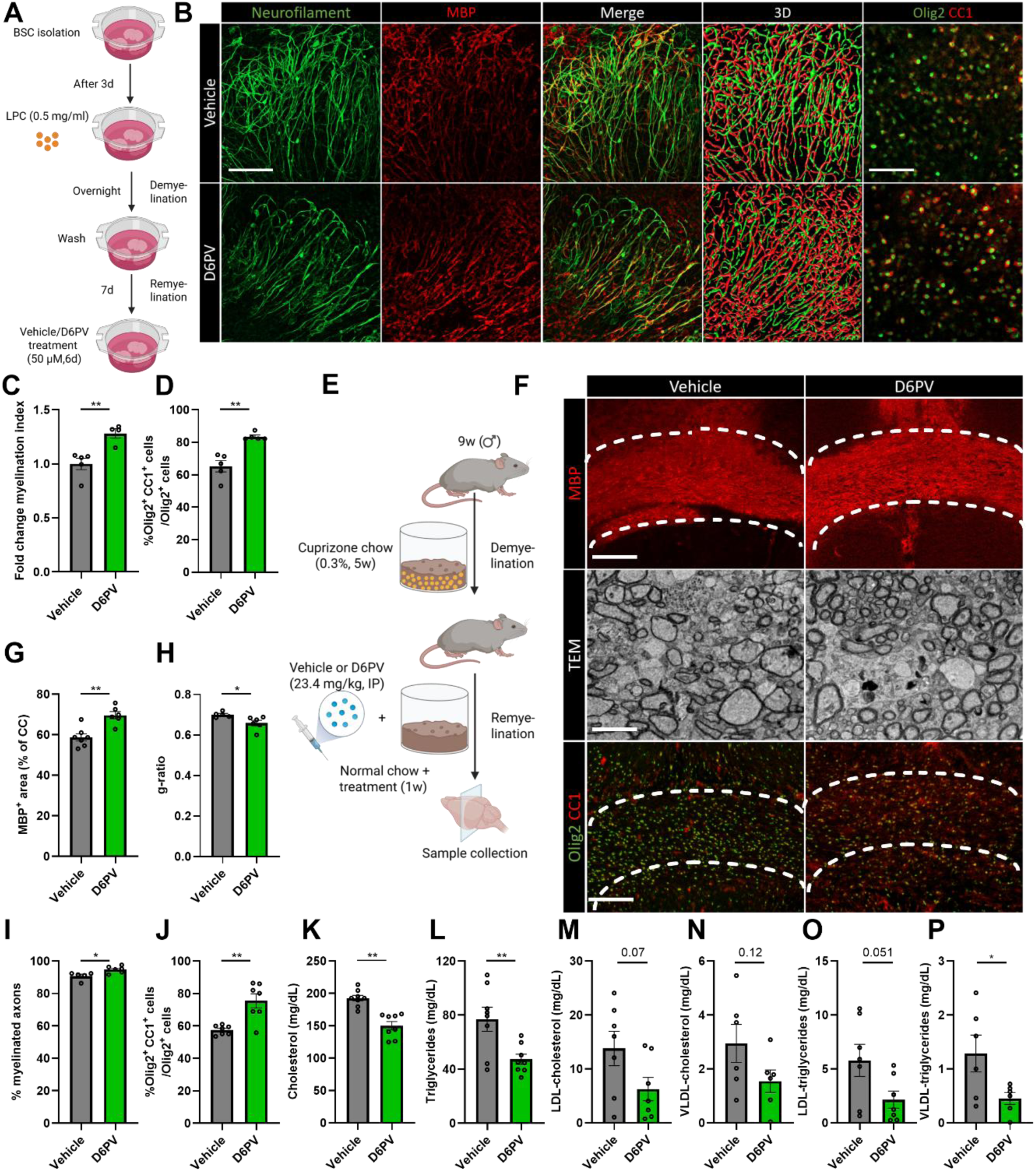
Peptide D6PV promotes remyelination in the cerebellar brain slice and cuprizone models. **A)** Schematic representation showing the experimental pipeline used to assess the impact of D6PV on remyelination in the *ex vivo* cerebellar brain slice model. **B**) Representative images and three-dimensional reconstruction of immunofluorescent MBP/NF and Olig2/CC1 stains of wild-type cerebellar brain slices exposed to vehicle or D6PV (50 µM). Scale bar, 100 µm. **C**) Relative number of MBP^+^ NF^+^ axons out of total NF^+^ axons in vehicle-treated and D6PV-treated cerebellar brain slices (n = 4-5 slices, 3 biological replicates). **D**) Percentage Olig2^+^ CC1^+^ cells within the Olig2^+^ cell population in vehicle-treated and peptide D6PV-treated cerebellar brain slices (n = 7 slices, 3 biological replicates). **E**) Schematic representation showing the experimental pipeline used to assess the impact of D6PV on remyelination in the cuprizone model. **F**) Representative images of immunofluorescent MBP and Olig2/CC1 stains and transmission electron microscopy (TEM) analysis of the corpus callosum (CC) from wild-type mice exposed to cuprizone, treated daily intraperitoneally (IP) with vehicle or D6PV (23.4 mg/kg) for 1 week during remyelination. The outer border of the CC is demarcated by a dotted line. Analysis was performed during remyelination. Scale bars, 200 µm (rows 1, 3) and 1 µm (row 2). **G**) Quantification of the MBP^+^ area of the CC from cuprizone mice treated with vehicle or D6PV during remyelination (n = 6-7 mice). **H,I**) Analysis of the g-ratio (the ratio of the inner axonal diameter to the total outer diameter, H, n = 7 mice) and percentage myelinated axon (I, n = 7 mice) in CC from cuprizone mice treated with vehicle or D6PV during remyelination. Quantification was performed on 250 axons/animal. **J**) Quantification of the percentage Olig2^+^ CC1^+^ cells out of total Olig2^+^ cells in the CC of cuprizone mice treated with vehicle or D6PV during remyelination (n = 7 mice). **K-P**) Concentrations of cholesterol (K), triglycerides (L), LDL-cholesterol (M), VLDL-cholesterol (N), LDL-triglycerides (O), and VLDL-triglycerides (P) in plasma of cuprizone mice treated with vehicle or D6PV. Data are represented as mean ± s.e.m. and statistically analyzed using the two-sided Student’s t-test. *, *p* < 0.05; **, *p* < 0.01. If *p* values are not indicated, the result was not significant.

To validate these findings *in vivo*, we employed the cuprizone model of toxic demyelination, which induces reproducible oligodendrocyte loss and myelin degradation in multiple brain areas, particularly the corpus callosum (CC) ^38,39^. Mice were fed 0.3% cuprizone chow for five weeks to induce demyelination, followed by one week of recovery during which D6PV was administered intraperitoneally on a daily basis (experimental design depicted in **Fig. 1E**). MBP immunostaining revealed a significant increase in myelin abundance in the CC of D6PV-treated mice compared to vehicle controls (**Fig. 1F,G**). Transmission electron microscopy confirmed these results, showing a reduction in the g-ratio (**Fig. 1F,H**), which represents the ratio of the inner axonal diameter to the total outer diameter, including the myelin sheath. A lower g-ratio reflects thicker myelin relative to axon size, indicating more efficient remyelination. Alongside improved axonal myelination, the percentage of myelinated axons was elevated in cuprizone mice exposed to D6PV (**Fig. 1F,I**), further supporting enhanced myelin regeneration. Consistent with the BSC model, D6PV-treated mice also exhibited an increased proportion of mature Olig2⁺ CC1⁺ oligodendrocytes within the CC (**Fig. 1F,J**), reinforcing the conclusion that D6PV facilitates oligodendrocyte maturation and functional remyelination. Notably, lipidomic profiling revealed that D6PV reduced both total and lipoprotein-associated serum cholesterol and triglyceride levels (**Fig. 1K-P**), underscoring its metabolic specificity. Together, these findings demonstrate that D6PV enhances remyelination across both *ex vivo* and *in vivo* models of chemically induced demyelination.

### D6PV does not alter EAE disease course or peripheral immune composition

To investigate the immunological impact of D6PV in the context of autoimmune-mediated neuroinflammation and demyelination, we employed the experimental autoimmune encephalomyelitis (EAE) model, a widely accepted preclinical paradigm for MS ^40^. Mice were immunized with the myelin oligodendrocyte glycoprotein peptide MOG_₃₅–₅₅_ and subsequently administered D6PV intraperitoneally at a dose of 23.4 mg/kg per day, beginning five days post-immunization (dpi; (experimental design depicted in **Fig. 2A**) - a strategic time point selected to coincide with the early priming phase of autoreactive T cells and the initiation of peripheral immune activation. Clinical disease scores were monitored longitudinally to capture the full trajectory of EAE progression, and immune profiling was performed at 28 dpi using multiparametric flow cytometry on cells isolated from both the spleen and inguinal lymph nodes.

**Figure 2:**
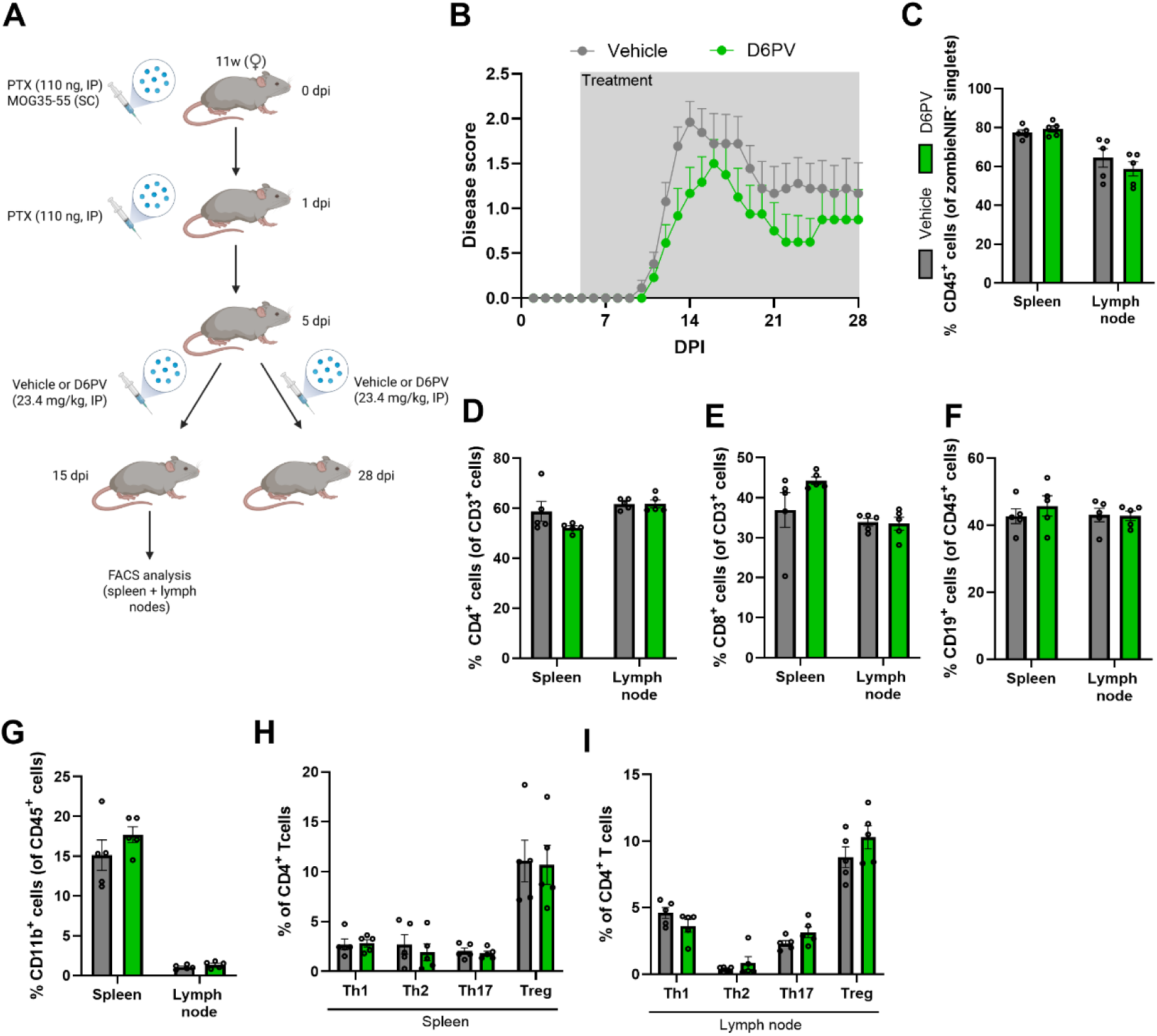
Peptide D6PV does not alter EAE disease score of peripheral immune composition. **A**) Schematic representation showing the experimental pipeline used to assess the impact of D6PV on disease score and peripheral immune composition in the experimental autoimmune encephalomyelitis (EAE) model. **B**) Disease score of 12w-old female wild-type mice in which EAE was induced. Starting 5 days post-immunization (dpi), animals were injected intraperitoneally (IP) with vehicle or D6PV (23.4 mg/kg; n = 15 mice) on a daily basis. **C**) Frequency of CD45^+^ cells in spleen and lymph nodes of EAE mice 15 dpi injected with vehicle or D6PV (n = 5 mice). **D-E**) Frequency of CD4^+^ (D) and CD8^+^ (E) cells within the CD3^+^ population in spleen and lymph nodes of EAE mice 15 dpi injected with vehicle or D6PV (n = 5 mice). **F-G**) Frequency of CD19^+^ (F) and CD11b^+^ (G) cells within the CD45^+^ population in spleen and lymph nodes of EAE mice 15 dpi injected with vehicle or D6PV (n = 5 mice). **H-I**) Frequency of T helper 1 (Th1), Th2, Th17 and regulatory T (Treg) cells within the naïve T cell population in spleen (H) and lymph nodes (I) of EAE mice 15 dpi injected with vehicle or D6PV (n = 5 mice). Data are represented as mean ± s.e.m. and statistically analyzed using two-sided two-way ANOVA. If *p* values are not indicated, the result was not significant.

D6PV-treated mice exhibited a modest attenuation in clinical severity compared to vehicle controls, however, this trend did not reach statistical significance, and no differences were observed in disease onset, peak neurological impairment, or recovery kinetics (**Fig. 2B**). These data suggest that D6PV does not substantially alter the course of EAE at the level of clinical symptoms. To further dissect potential immunological effects, we quantified major immune cell populations across lymphoid compartments (gating strategy, see **Suppl. Fig. S1**) 15 dpi. The frequencies of CD45⁺ hematopoietic cells (**Fig. 2C**), CD4⁺ and CD8⁺ T cells (**Fig. 2D-E**), CD19⁺ B cells (**Fig. 2F**), and CD11⁺ myeloid cells (**Fig. 2G**) were comparable in the spleen and inguinal lymph nodes of D6PV- and vehicle-treated animals, indicating no gross perturbation of lymphocyte or myeloid cell homeostasis. Within the CD4⁺ T cell compartment, we further assessed canonical effector subsets associated with EAE pathogenesis. The proportions of Th1 (IFNγ⁺), Th2 (IL-4⁺), and Th17 (IL-17⁺) cells - key drivers of neuroinflammation - were indistinguishable between groups in both the spleen (**Fig. 2H**) and inguinal lymph nodes (**Fig. 2I**). Similarly, the abundance of FOXP3⁺ regulatory T cells, which are known to reduce EAE disease severity, remained stable in frequency in both lymphoid tissues (**Fig. 2H-I**). Taken together, these findings suggest that D6PV does not significantly impact EAE disease severity or alter systemic immune architecture in the EAE model. These findings further suggest that the pro-regenerative effect of D6PV observed in the cuprizone model is likely driven by local CNS mechanisms rather than modulation of peripheral immunity.

**Supplementary Figure 1:**
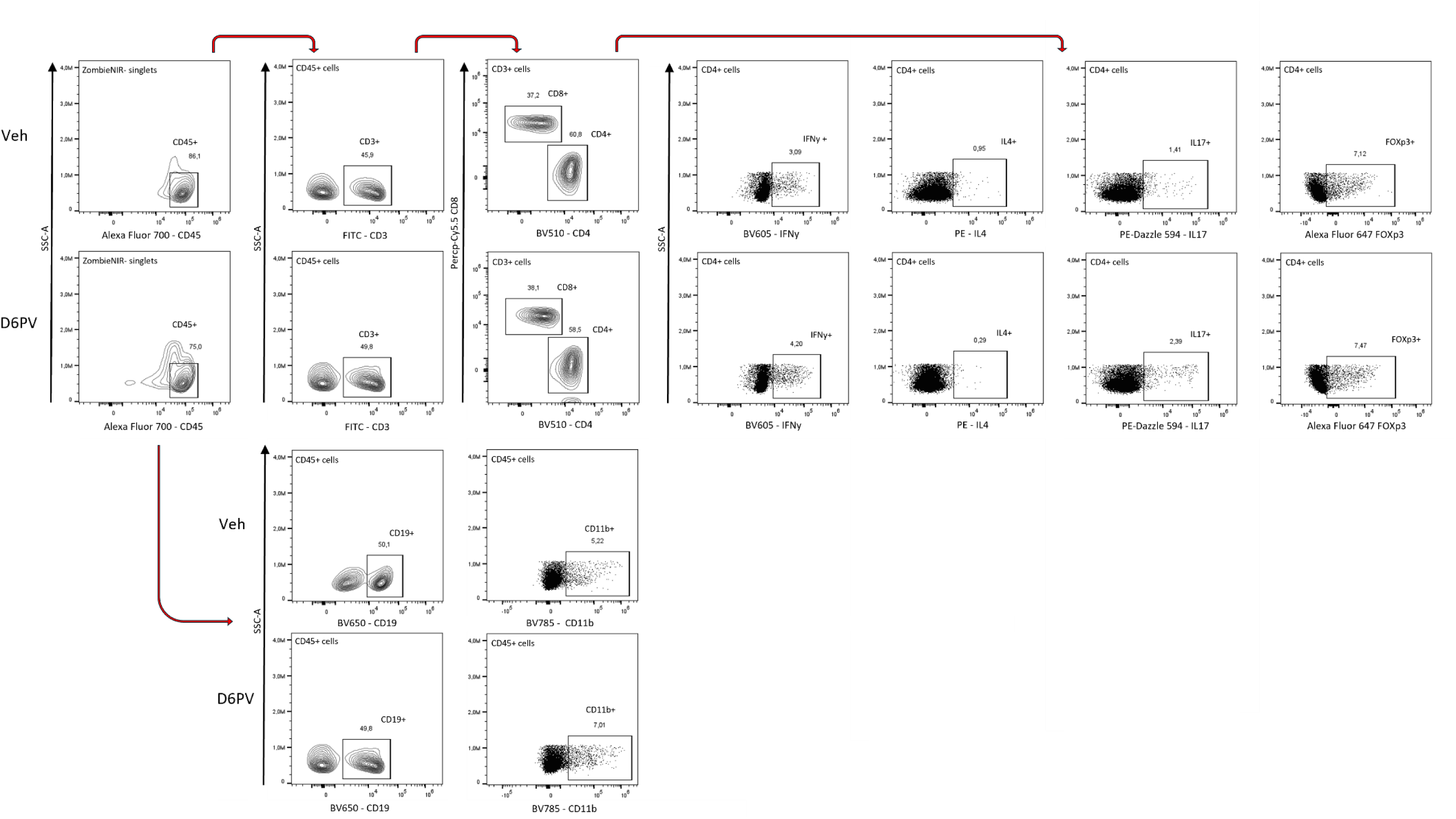
Gating strategy of FACS EAE immune panel. Representative flow cytometric plots are shown.

### D6PV does not impact the metabolic, inflammatory, and regenerative properties of foam cells

We and others have recently demonstrated that sustained intracellular accumulation of myelin-derived lipids, and their sequestration into lipid droplets, drives phagocytes toward a pro-inflammatory state that impairs remyelination in both the BSC and cuprizone models ^11–13^. Given that D6PV is a potent activator of LPL, facilitating the hydrolysis and clearance of triglycerides ^20^, the predominant lipid species stored within lipid droplets ^41^, we hypothesized that D6PV might enhance remyelination by reducing the lipid droplet burden in myelin-laden phagocytes and thereby biasing these foam cells toward a pro-regenerative phenotype. To assess whether D6PV modulates phagocyte lipid metabolism or phenotype, primary bone marrow-derived macrophages (BMDMs) were exposed to isolated myelin (24h or 72h) in the presence or absence of D6PV (10 µM). D6PV treatment did not alter the lipid droplet load, as indicated by unchanged BODIPY fluorescence and Oil Red O (ORO) staining in both control and myelin-exposed BMDMs (**Fig. 3A-C**). Likewise, demyelinated BSCs exposed to D6PV showed no difference in the frequency of F4/80⁺ BODIPY⁺ microglia (**Fig. 3D,E**). Consistent with these observations, intracellular levels of esterified cholesterol - a major determinant of lipid droplet formation - as well as total and free cholesterol, were unaffected by D6PV (**Fig. 3F**). D6PV also failed to promote cholesterol efflux via ABCA1 (**Fig. 3G**), a key pathway mediating lipid recycling in the CNS ^11,14,37^. Surprisingly, although D6PV did not reduce intracellular lipid stores, phagocytosis assays revealed a modest increase in uptake of pHrodo-labeled myelin following D6PV exposure (**Fig. 3H**), suggesting enhanced engulfment capacity.

**Figure 3:**
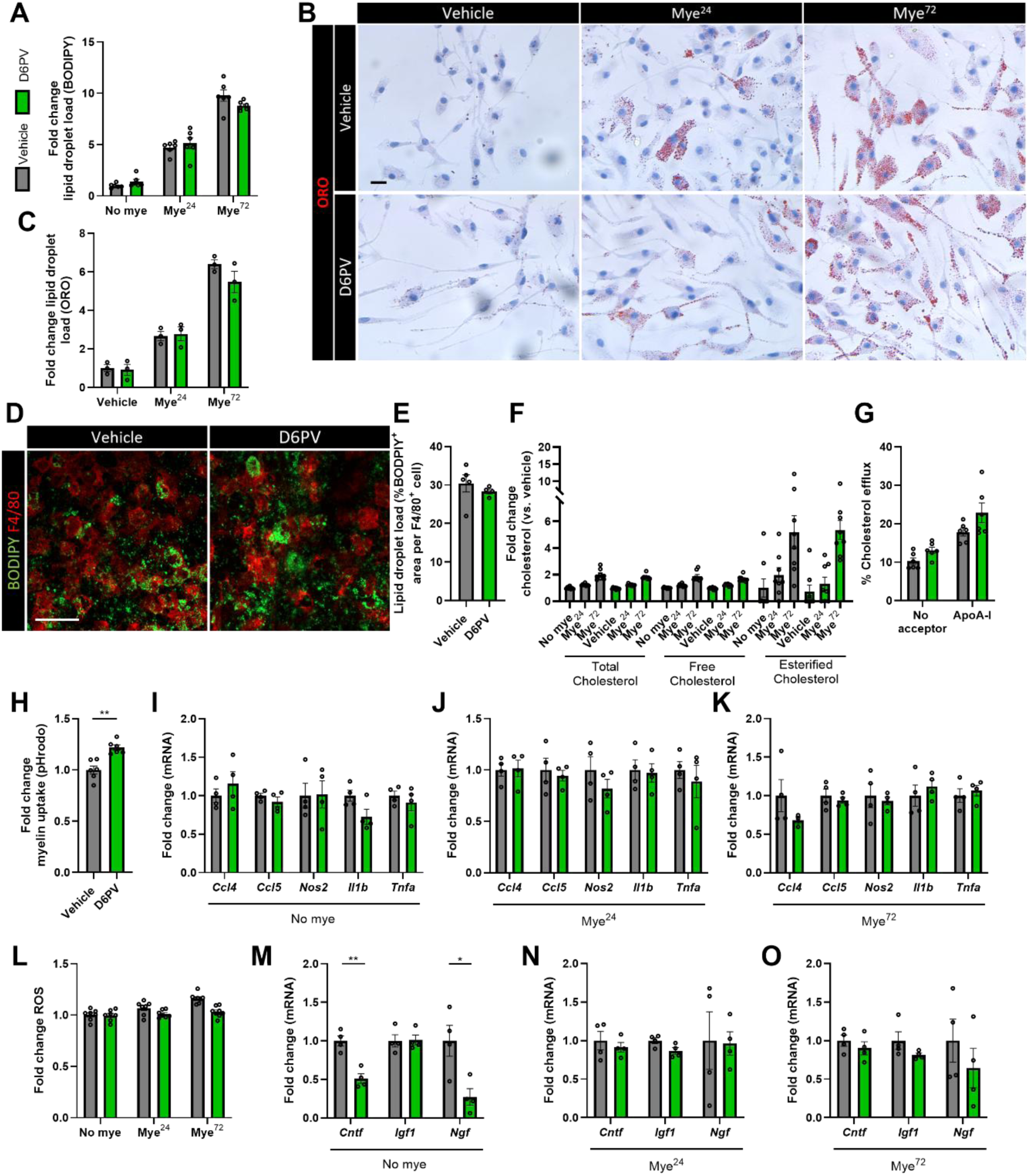
Peptide D6PV does not affect the metabolic, inflammatory, or regenerative phenotype of foam cells. **A**) Mean fluorescence intensity of BODIPY in wild-type bone marrow-derived macrophages (BMDMs) treated with vehicle or myelin (100 µg/ml), with or without D6PV (10 µM) for 24 or 72h (n = 6 cultures, 3 biological replicates). **B-C**) Representative images (B) and quantification (C) of Oil red O (ORO) staining of wild-type BMDMs treated with vehicle or myelin, with or without D6PV for 24 or 72h. Lipid droplet load is quantified as percentage ORO^+^ area per cell (n = 3 cultures, 3 biological replicates). Scale bar, 50 µm. **D-E**) Representative images (D) and quantification (E) of BODIPY/F4/80 stain of wild-type cerebellar brain slices treated with vehicle or peptide D6PV (50 µM). Lipid droplet load was quantified as percentage of BODIPY^+^ area per F4/80^+^ cell (n = 4-5 slices, 2 biological replicates). Scale bar, 100 µm. **F**) Quantification of total cholesterol, free cholesterol, and esterified cholesterol in wild-type BMDMs treated with vehicle or myelin, with or without D6PV for 24 or 72h. Data is depicted as fold change vs. vehicle-treated BMDMs without D6PV (n = 8 cultures, 2 biological replicates). **G**) Relative cholesterol efflux capacity of wild-type BMDMs loaded with myelin and treated with or without D6PV for 24h, without or with ApoA-I mimetic peptide 5A (50 µg/ml, 4h) as cholesterol acceptor (n = 6 cultures, 2 biological replicates). **H**) Geomean of pHrodo in wild-type BMDMs treated with vehicle or D6PV for 24h, followed by 1.5h exposure to pHrodo-labeled myelin (n = 6 cultures, 2 biological replicates). **I-K)** mRNA expression of *Ccl4*, *Ccl5*, *Nos2*, *Il1b*, and *Tnfa* in wild-type BMDMs treated with vehicle or myelin, with or without D6PV for 24 or 72h, followed by LPS/IFNγ (100 ng/mL) stimulation for 6h (n = 4 cultures, 2 biological replicates). **L**) ROS production in wild-type BMDMs treated with vehicle or myelin, with or without D6PV for 24 or 72h, followed by PMA (100 ng/mL) stimulation for 15 minutes (n = 7-8 cultures, 2 biological replicates). **M-O**) mRNA expression of *Cntf*, *Igf1*, and *Ngf* in wild-type BMDMs treated with vehicle or myelin, with or without D6PV for 24 or 72h (n = 4 cultures, 2 biological replicates). Data are represented as mean ± s.e.m. and statistically analyzed using the two-sided Student’s t-test (A, C, E, G-O) or the two-sided two-way ANOVA (F). *, *p* < 0.05; **, *p* < 0.01. If *p* values are not indicated, the result was not significant.

Beyond its minimal effects on foam cell formation and lipid handling, D6PV did not alter the inflammatory phenotype of either control or myelin-laden macrophages, as expression of *Ccl4*, *Ccl5*, *Nos2*, *Il1b*, and *Tnfa* remained unchanged in activated BMDMs under both conditions (**Fig. 3I-K**), and reactive oxygen species production was similarly unaffected (**Fig. 3L**). Moreover, analysis of neurotrophic factors demonstrated that D6PV did not increase *Cntf*, *Igf1*, and *Ngf* expression in control and myelin-exposed BMDMs (**Fig. 3M-O**). In fact, control BMDMs displayed a reduced expression of *Cntf* and *Ngf* upon D6PV exposure (**Fig. 3M**). Together, these results indicate that D6PV does not substantially reshape the metabolic, inflammatory, or pro-regenerative phenotype of control or myelin-loaded macrophages *in vitro*.

### D6PV promotes oligodendrocyte precursor cell maturation

So far, we have found that D6PV promotes remyelination both *ex vivo* and *in vivo*, but exerts minimal effects on (foamy) phagocytes *in vitro*. To firmly establish whether phagocytes are dispensable for their pro-remyelinating activity, we depleted microglia in BSCs using a clodronate liposome-based approach previously established by our research group ^15,16,37^. D6PV enhanced remyelination in BSCs exposed to empty liposomes (**Fig. 4A,B**), consistent with our earlier findings (**Fig. 1B,C**), and retained this effect in BSCs depleted of microglia using clodronate-loaded liposomes (**Fig. 4A,B**). Correspondingly, an increased proportion of mature oligodendrocytes was observed in BSCs exposed to both empty or clodronate-loaded liposomes (**Fig. 4A,C**). These data indicate that D6PV promotes remyelination in a phagocyte-independent manner.

**Figure 4:**
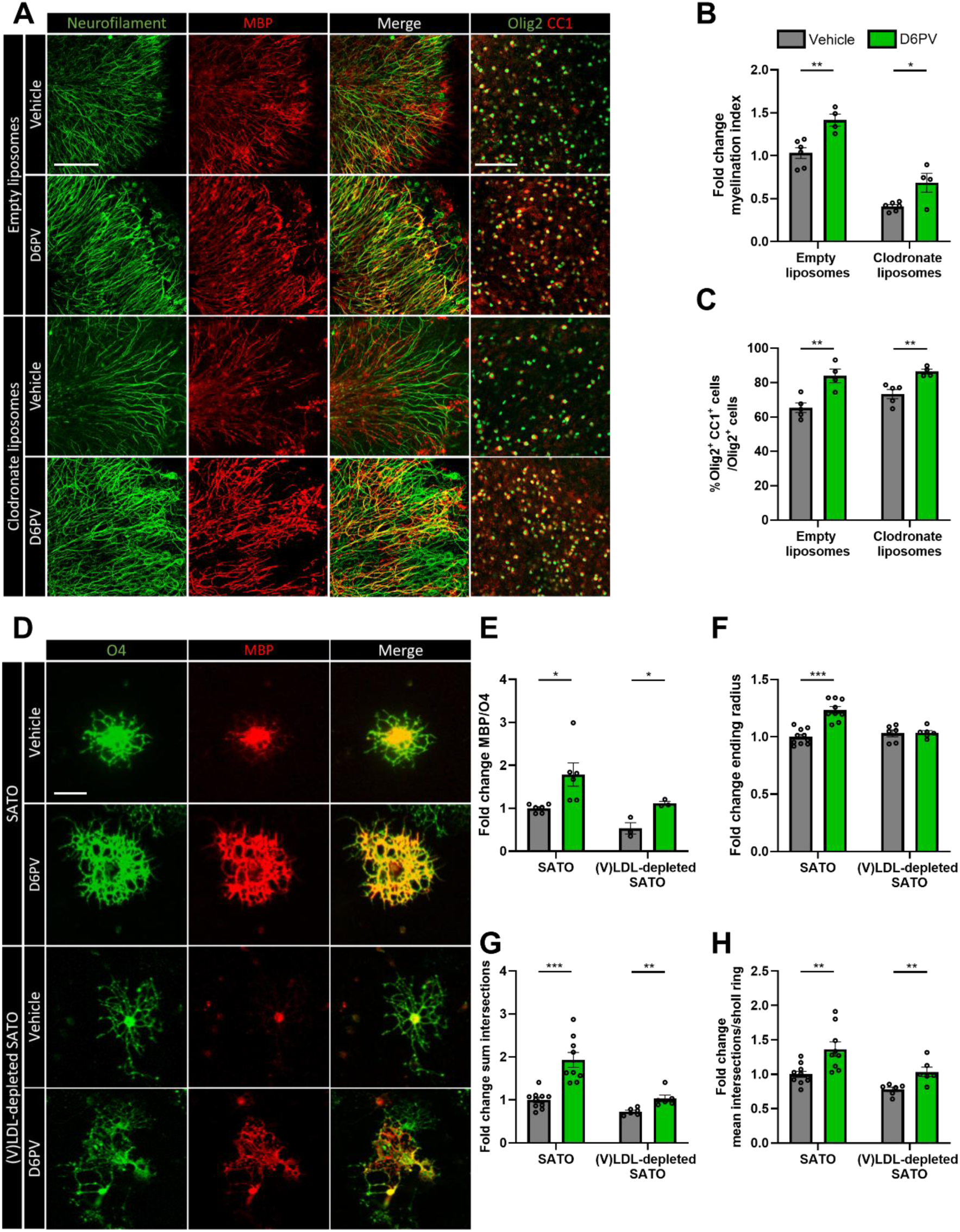
Peptide D6PV improves oligodendrocyte precursor cell maturation. **A**) Representative images of immunofluorescent MBP/NF and Olig2/CC1 stains of wild-type cerebellar brain slices exposed to empty or clodronate liposomes (0.5 mg/mL) and exposed to vehicle or D6PV (50 µM). Scale bar, 100 µm. **B**) Fold change of relative number of MBP^+^ NF^+^ axons out of total NF^+^ axons in cerebellar brain slices treated with empty or clodronate liposomes and with vehicle or D6PV (n = 4-6 slices, 2 biological replicates). **C**) Percentage Olig2^+^ CC1^+^ cells within the Olig2^+^ cell population in cerebellar brain slices treated with empty or clodronate liposomes and with vehicle or D6PV (n = 4-5 slices, 2 biological replicates). **D-H**) Representative immunofluorescent images (D) and quantifications (E-H) of wild-type oligodendrocyte precursor cells (OPCs) exposed to a regular or (very) low-density lipoprotein ((V)LDL)-depleted differentiation cocktail for 6 days and treated daily with vehicle or D6PV (10 µM). OPCs were stained for MBP (mature OLNs) and O4 (committed OLNs), and differentiation was quantified by measuring the MBP/O4 ratio (E, n = 3-6 cultures, 3 biological replicates) or applying Sholl analysis (sum intersections, mean intersections/sholl ring and ending radius; F-H, n = 6-10 cultures, 3 biological replicates). Scale bar, 25 µm. Data are represented as mean ± s.e.m. and statistically analyzed using the two-sided Student’s t-test. *, *p* < 0.05; **, *p* < 0.01; ***, *p* < 0.001. If *p* values are not indicated, the result was not significant.

Based on these findings, we assessed the direct impact of D6PV on OPC differentiation. To this end, primary mouse OPCs were exposed to a differentiation cocktail and D6PV for 6 days. Fluorescence analysis revealed a significant increase in the ratio of the mature OL marker MBP to the pre-OL marker O4 (**Fig. 4D,E**), indicating enhanced OPC–OLN maturation. Sholl analysis further confirmed increased dendritic complexity and greater terminal branching radius in D6PV-treated cells (**Fig. 4F-H**), consistent with the generation of highly ramified, myelination-competent OLNs. Given ApoC-II/III’s canonical role in activating LPL and regulating triglyceride-rich lipoprotein metabolism, we next sought the determine whether exogenous lipoproteins in the culture medium were required for D6PV’s promyelinating activity. We observed that D6PV retained its ability to enhance the ratio of the mature OLN marker MBP over the pre-OLN marker O4 in lipoprotein-depleted medium (**Fig. 4D,E**). As expected, lipoprotein depletion moderately impaired OPC maturation overall (**Fig. 4D,E**). Alongside enhancing the MBP/O4 ratio, D6PC continued to increase the number and complexity of dendritic branches in lipoprotein-depleted medium (**Fig. 4F-H**). Collectively, these findings indicate that D6PV promotes OPC maturation and OLN formation independently of lipoprotein hydrolysis or exogenous lipoprotein availability, establishing a phagocyte-independent, direct promyelinating mechanism.

### D6PV enhances OPC maturation by enhancing oxidative phosphorylation

To delineate molecular programs engaged by D6PV, we performed bulk RNA sequencing of OPCs. Differential expression analysis identified 2095 genes that distinguished untreated and D6PV-exposed OPCs, with 734 genes upregulated and 1361 genes downregulated following D6PV treatment (**Fig. 5A**). Pathway enrichment analysis using Ingenuity Pathway Analysis (IPA) revealed a pronounced enrichment of mitochondrial metabolic pathways, including oxidative phosphorylation (z-score 6.782), respiratory electron transport (z-score 6.403), mitochondrial translation (z-score 5.385), complex I biogenesis (z-score 4.690), and mitochondrial protein degradation (z-score 4.536), together with a marked suppression of mitochondrial dysfunction (z-score −5.080) (**Fig. 5B**). These transcriptional changes suggest that D6PV induces coordinated adaptations in mitochondrial physiology, consistent with enhanced oxidative phosphorylation and fatty acid β-oxidation in OPCs. Upstream regulator analysis further indicated reduced activation of inflammatory signaling pathways (e.g., CXCR4, SOD2, MAP2K4, SP1, and LPS), alongside increased engagement of lipid-responsive and mitochondrial-associated regulators, most notably components of the PPAR signaling axis (e.g., PPARGC1A, PPARGC1B, PPARA, NRF1) and the AMPK signaling axis (e.g., PRKAA1, PRKAA2, STK11, CAB39L) (**Fig. 5C**). Collectively, these data support the conclusion that D6PV facilitates OPC maturation by modulating metabolic and inflammatory transcriptional programs in a manner that favors mitochondrial oxidative metabolism and fatty acid utilization in OPCs.

**Figure 5:**
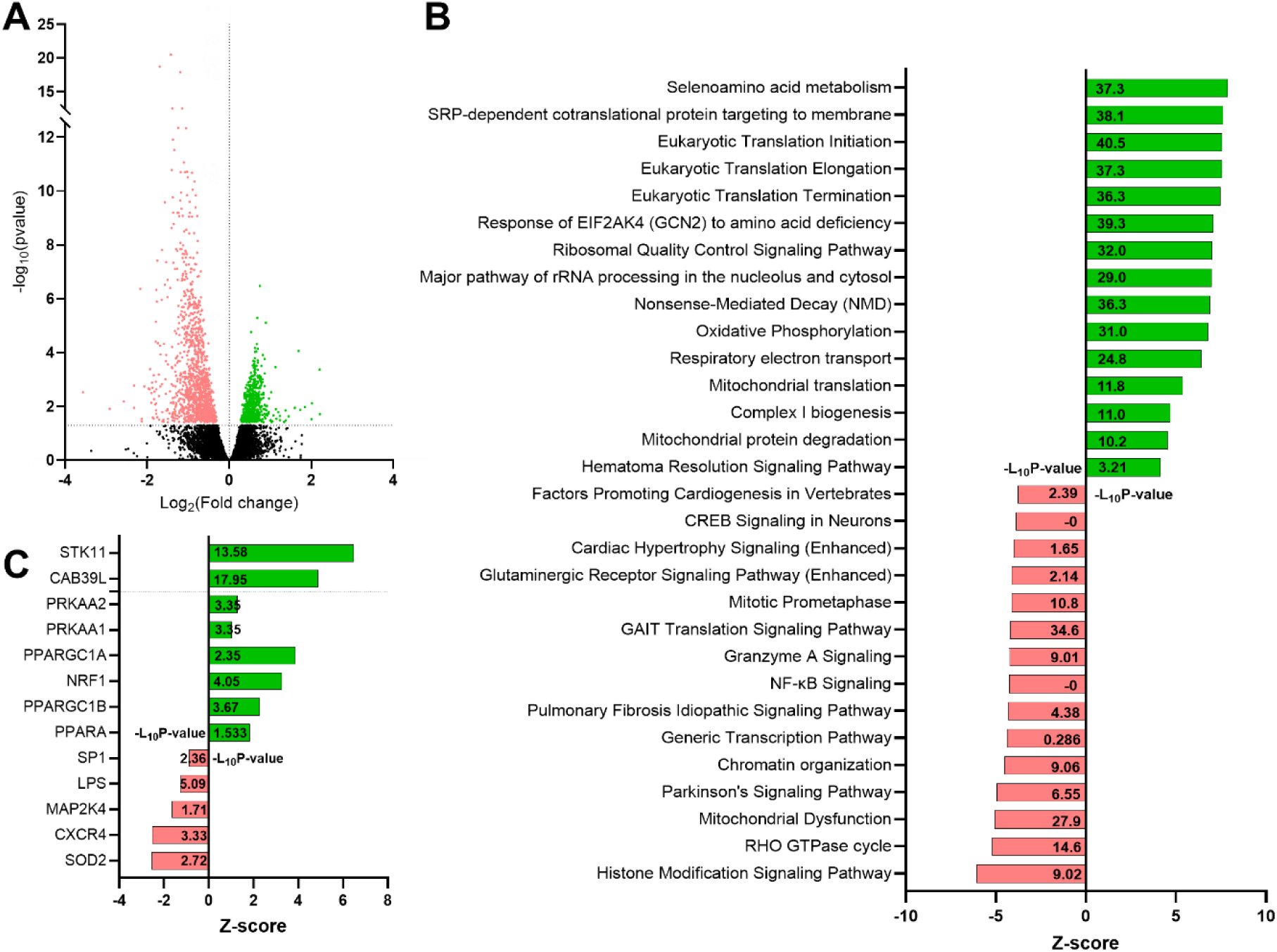
Peptide D6PV promotes mitochondrial metabolic pathways in oligodendrocyte precursor cells. **A**) Bulk RNA sequencing was performed on oligodendrocyte precursor cells (OPCs) exposed to vehicle or D6PV (10 µM) for 24h. Differentially expressed genes were used as input for core analysis in Ingenuity Pathway Analysis (IPA) (n = 5 cultures, cut-off criteria p < 0.05). **B**) Pathway analysis showing up- and downregulated canonical pathways of D6PV-treated OPCs. – Log_10_(Pvalue) and down-or upregulated canonical pathways with corresponding Z-scores are indicated at y- and x-axis, respectively. **C**) Upstream analysis showing down- and upregulated regulators in D6PV-treated OPCs.

## Discussion

Accumulating evidence underscores the essential role of lipid metabolic pathways in orchestrating OPC maturation and remyelination ^42,43^. In this context, changes in ApoC-II, ApoC-III, LPL abundance, and lipid profiles characteristic of ApoC-II/ApoC-III imbalances, are closely associated with MS disease progression ^17–19,23–25^. In this study, we identify the dual ApoC-II mimetic–ApoC-III antagonist peptide D6PV as a previously unrecognized regulator of OPC metabolic state that promotes their differentiation and enhances remyelination in the CNS. We show that D6PV enhances mitochondrial oxidative phosphorylation, thereby boosting fatty acid β-oxidation, suppressing inflammatory transcriptional programs, and enabling OPCs to progress toward a mature, myelinating phenotype. Together, these findings uncover a metabolism-linked mechanism that promotes CNS regeneration and position D6PV as a promising therapeutic strategy to enhance remyelination.

By using *in vitro*, *ex vivo*, and *in vivo* preclinical models, we provide evidence that D6PV promotes remyelination by inducing a metabolically favorable phenotype that supports OPC differentiation. Although ApoC-II/III classically modulate LPL-mediated triglyceride hydrolysis, this pro-differentiation effect occurred independently of lipoproteins *in vitro*, suggesting alternative molecular roles for ApoC-II/III. In light of the latter, while literature on the direct impact of ApoC-II on cell biology is scarce, ApoC-III is increasingly recognized as a potent driver of lipid-induced inflammation and tissue pathology. For instance, in chronic kidney disease, ApoC-III directly promotes inflammation by activating the NLRP3 inflammasome in human monocytes, thereby amplifying systemic inflammatory signaling and contributing to organ injury ^44^. Mechanistic studies further show that ApoC-III induces endothelial activation, increases TNF-α expression, and disrupts tight junction integrity, ultimately driving vascular dysfunction ^45^. Additionally, ApoC-III promotes inflammation-dependent neovascularization, implicating it in maladaptive tissue remodeling ^46^. Consistent with these findings, our RNA sequencing analysis revealed that D6PV-exposed OPCs exhibited reduced activation of inflammatory signaling pathways. Together, these results establish a pro-regenerative capacity of D6PV in the CNS, although whether this effect is predominantly mediated by ApoC-II–mimetic activity, ApoC-III antagonism, or a combination of both remains to be determined.

We and others have recently identified foam cell formation as a critical bottleneck to efficient remyelination ^11,12^. Excessive uptake of myelin-derived lipids during demyelination drives the accumulation of lipid droplet–engorged macrophages and microglia that adopt a highly inflammatory, repair-inhibitory phenotype ^11^. Disrupted lipid uptake, intracellular remodeling, and efflux underlie the emergence of this maladaptive state ^16,47,48^. Despite triglycerides being a major content of lipid droplets and previous studies showing that LPL controls myeloid cell lipid metabolism and function ^17,49^, our data indicate that D6PV does not substantially reprogram lipid metabolic pathways in foamy macrophages. This distinction could highlight a cell type-specific mechanism, wherein D6PV directly enhances the metabolic and inflammatory state of OPCs to support their differentiation and remyelination, rather than acting indirectly by modulating phagocyte lipid metabolism. Future experiments using cell type-specific genetic or pharmacological approaches could determine why OPCs are selectively responsive to D6PV and elucidate the precise metabolic and signaling pathways underlying this pro-regenerative effect.

Myelin synthesis imposes a substantial energetic burden on developing OLNs, but this investment ultimately reduces axonal energy demand and accelerates signal conduction ^50^. Although glucose serves as the primary metabolic fuel, myelinating OLNs also utilize lactate to generate ATP and to supply carbon for myelin lipid synthesis, and lactate availability promotes OPC proliferation, differentiation, and myelination, even under hypoglycemic conditions ^51–54^. Consistent with these requirements, OPCs and differentiating OLNs exhibit a high rate of mitochondrial oxidative phosphorylation (OXPHOS) and contain an abundance of long, tubular mitochondria ^55^, a morphology associated with elevated respiratory capacity ^56^. As OLNs mature, mitochondrial density decreases, and organelles adopt a more fragmented architecture, suggesting a metabolic shift toward reduced reliance on OXPHOS, a phenomenon observed across multiple differentiating cell types ^57,58^. Despite this shift, OLNs remain metabolically flexible. When glucose levels decrease, OLNs can switch from glycolysis to fatty acid β-oxidation, using fatty acids derived from peripheral lipid stores or from autophagy-mediated recycling of myelin lipids to sustain ATP production for OXPHOS ^59,60^. The importance of this metabolic adaptability is underscored in developmental and disease contexts. In intrauterine growth restriction, impaired mitochondrial function, marked by downregulation of OXPHOS and electron transport chain genes, reduced mitochondrial respiration, and diminished ATP production, leads to an early arrest in OPC differentiation ^61^. Conversely, enhancing mitochondrial competence can promote OLN survival and differentiation, as demonstrated in Alzheimer disease-related models where increased expression of adenylate kinase 5 enhances AMPK signaling, stimulates OXPHOS, and suppresses inflammation and apoptosis in OLNs ^62^. In line with the central role of mitochondrial activity in lineage progression, our data now demonstrate that D6PV treatment increases fatty acid β-oxidation and OXPHOS in differentiating OPCs, further supporting the concept that enhancing mitochondrial respiration can facilitate OLN maturation.

Our findings demonstrate that ApoC-II mimetic peptide D6PV robustly enhances OPC differentiation and accelerates remyelination. Notably, however, D6PV did not attenuate inflammatory responses *in vitro*, as BMDMs cultured with D6PV showed no reduction in pro-inflammatory markers, nor did they display increased expression of neurotrophic mediators, consistent with previous observations in myelin-laden macrophages ^11,12,63,64^. On that same note, D6PV treatment did not ameliorate clinical disease severity in EAE, nor did it reduce the proportion of pro-inflammatory immune cells infiltrating the CNS. Together, these results indicate that the pro-regenerative actions of D6PV operate independently of modulation of the neuroinflammatory environment. Rather, its therapeutic benefit appears limited to the promotion of remyelination. This implies that D6PV alone is unlikely to provide clinical benefit in disorders dominated by chronic neuroinflammation. Thus, leveraging its full therapeutic potential will likely require combination therapy, pairing D6PV with agents that effectively dampen neuroinflammation.

In conclusion, our findings position metabolic reprogramming of OPCs as a key determinant of successful remyelination and establish the dual ApoC-II mimetic/ApoC-III antagonist peptide D6PV as a previously unrecognized pro-regenerative regulator of this process. By selectively enhancing mitochondrial oxidative metabolism and fatty acid utilization in OPCs, without broadly suppressing neuroinflammation, D6PV reveals a cell-intrinsic metabolic constraint that can be therapeutically targeted to promote myelin repair. These results advance the emerging concept that remyelination failure is not solely driven by inflammatory burden or limited progenitor availability, but also suggests that modulation of OPC metabolism can be used as a strategy to promote remyelination. Elucidating the precise modulator targets of D6PV and defining its optimal integration with anti-inflammatory therapies will be essential toward translating metabolic interventions into effective remyelination strategies for MS and related demyelinating disorders.

## Acknowledgements

We thank L. Timmermans, M.P. Tulleners, W. Vandendries, and M. Jans for excellent technical assistance. BM is funded by the Research Foundation Flanders (FWO Vlaanderen; 11PAO24N). LB is funded by the Research Foundation Flanders (FWO Vlaanderen; 1SH6G24N). KP is funded by the Research Foundation Flanders (FWO Vlaanderen; 1104425N). KK is funded by the Research Foundation Flanders (FWO Vlaanderen; 11PLR24N). SGSV is funded by the Research Foundation Flanders (FWO Vlaanderen; 12B1I24N). ML is funded by the Research Foundation Flanders (FWO Vlaanderen; 1210026N). JJAH is funded by the Research Foundation Flanders (FWO Vlaanderen; G0A7922, G0A7922, S01623N) and the Charcot Research Foundation (CHARCO22HJ, CHARCO23HJ, CHARCO24HJ). JFJB is funded by the Research Foundation Flanders (FWO Vlaanderen; G075823, G0A3B24), Charcot Research Foundation (CHARCO23BJ, CHARCO24BJ,CHARCO25BJ), Geneeskundige Stichting Koningin Elisabeth (G.S.K.E; GSKE-BOGJ), MS Liga Vlaanderen (MSLIGABOGJ) and the special research fund Hasselt University (22DOC38BOF, 23INC06BOF). SV is funded by the Research Foundation Flanders (FWO Vlaanderen; 1245724N) and Charcot Research Foundation (CHARCO24VS). We acknowledge the Advanced Optical Microscopy Centre at Hasselt University for support with confocal microscopy experiments, which were made possibly by the Research Foundation Flanders (FWO Vlaanderen; G0H3716N, I001222N). The funding agencies had no role in the design, analysis, or writing of the article.

## Author contributions

BM, JFJB, and SV conceived experiments. BM, LB, KP, FW, KK, JG, SGSV, YJ, ML, and SV performed experiments. BM and SV analyzed data. BM, LB, KP, FW, KK, JG, SGSV, EW, YJ, YD, TV, TD, EG, MR, ATR, ML, JJAH, JFJB, and SV discussed results. BM, JFJB, and SV wrote the manuscript. BM, LB, KP, FW, KK, JG, SGSV, EW, YJ, YD, TV, TD, EG, MR, ATR, ML, JJAH, JFJB, and SV revised the manuscript.

