## Supplementary figures and images for "The ApoC2 mimetic peptide D6PV enhances remyelination by stimulating oxidative phosphorylation in oligodendrocytes"

### Supplementary Figure 1

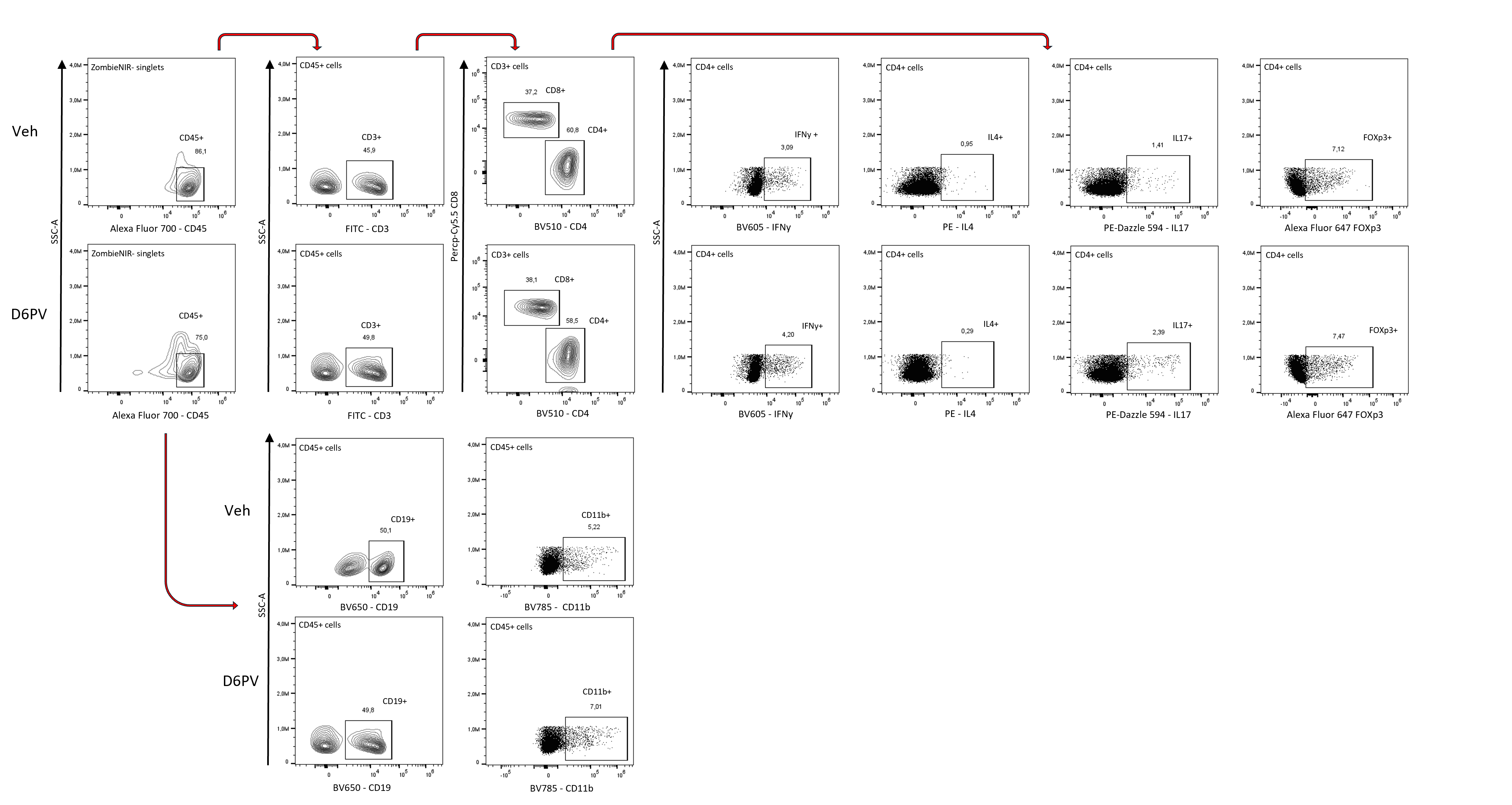
